# Episodic memory precision and reality monitoring following stimulation of angular gyrus

**DOI:** 10.1101/2021.09.15.460461

**Authors:** S. Kwon, F.R. Richter, M. J. Siena, J.S. Simons

## Abstract

The qualities of remembered experiences are often used to inform ‘reality monitoring’ judgments, our ability to distinguish real and imagined events (Johnson & Raye, 1981). Previous experiments have tended to investigate only whether reality monitoring decisions are accurate or not, providing little insight into the extent to which reality monitoring may be affected by qualities of the underlying mnemonic representations. We used a continuous-response memory precision task to measure the quality of remembered experiences that underlie two different types of reality monitoring decisions: agency decisions that distinguish actions performed by participants and the experimenter, and perceptual decisions that distinguish perceived and imagined experiences. The data revealed memory precision to be associated with higher accuracy in both agency and perceptual reality monitoring decisions, with reduced precision linked with a tendency to misattribute self-generated experiences to external sources. We then sought to investigate the possible neurocognitive basis of these observed associations by applying brain stimulation to a region that has been implicated in precise recollection of personal events, left angular gyrus. Stimulation of angular gyrus selectively reduced the association between memory precision and self-referential reality monitoring decisions, relative to control site stimulation. Angular gyrus may, therefore, be important for the ability to imbue remembered experiences with a sense of self-agency, a key component of ‘autonoetic consciousness’ that characterises episodic memory (Tulving, 1985).

## Introduction

Reality monitoring refers to a rememberer’s ability to keep track of whether their memories are veridical, as opposed to products of their imagination (Johnson & Raye, 1981). To explain how rememberers may keep a grasp on reality, a large body of previous research on reality monitoring has supported a prominent theoretical account: the Source Monitoring Framework (Johnson et al., 1993). The key hypothesis of this framework is that reality monitoring involves considering the features of retrieved memories against characteristics expected of real and imaginary experiences. For example, memories full of vivid visuoperceptual detail might be more likely to be real than those primarily comprising internally generated thoughts (Johnson et al., 1988). One prediction of this framework is that reality monitoring ability may depend, at least in part, on the precision with which details of past experiences are remembered (Simons et al., 2021; Simons et al., 2017). If, for example, a memory is relatively vague and imprecise, the rememberer may struggle to distinguish whether it relates to an event that actually occurred, or one that might have been imagined. Previous experiments on reality monitoring have tended to focus only on whether reality monitoring decisions are accurate or not, leaving unresolved the degree to which reality monitoring decisions are affected by the quality of the underlying memories. We aimed to bridge previously separate areas of research relating to reality monitoring and memory precision, by asking the following research questions: (1) to what extent does reality monitoring ability depend on the precision with which memories are retrieved? and (2) what is the neurocognitive basis of this possible dependency?

These research questions have implications for the understanding of symptoms concerning psychosis, such as hallucinations. It is possible that in schizophrenia, for example, patients’ memories lack precision, obscuring the self-referential characteristics that distinguish them as personal memories (Bentall et al., 1991; Seal et al., 1997; Waters et al., 2006; Woodward et al., 2007). Patients may therefore misattribute their past experiences or actions to another person, exhibiting an ‘It had to be you’ effect or ‘agency externalisation bias’ (Johnson et al., 1981; Johnson & Raye, 1981).

Conversely, imagined experiences full of unusually vivid or precise details may resemble perceptual features often expected of veridical memories. Patients with schizophrenia may therefore experience them as hallucinations, possibly thinking: ‘It’s so vivid, it had to be real’ (Mondino et al., 2019). The patients may, for example, report that they had seen objects move even when, in fact, they had imagined the object, exhibiting a ‘perceptual externalisation bias’ (Johnson & Raye, 1981).

Hallucinations are not only a symptom of mental illness. Previous studies have observed ‘non-clinical voice hearers’ to report experiences of hallucinations despite no clinical diagnosis or need for care (Baumeister et al., 2017). Such healthy people who are prone to hallucinations may misattribute imagined stimuli as real, for example, exhibiting externalisation bias. Previous studies involving large samples of healthy people sought to test this possible link between proneness to experience hallucination and externalisation bias, and some of these studies have indeed observed such correspondence (Allen et al., 2006; Collignon et al., 2005; Larøi et al., 2004), whereas other studies have not (Alderson-Day et al., 2019; Aynsworth et al., 2017; Garrison et al., 2017; Moseley et al., 2021). Note, however, that all these studies have used categorical responses to measure their participants’ mental experiences, which might have obscured variability in the qualities of those experiences.

To devise a more sensitive measure of remembered experiences, the current study combined reality monitoring tasks with a continuous-response long-term memory precision task adapted from the working memory literature (Bays et al., 2009; Richter et al., 2016). Richter, Cooper, and colleagues (2016) asked their participants to first study a series of displays consisting of objects overlaid at random locations around a circle and, in a later memory test phase, recreate the exact location of each object using a continuous response dial. We incorporated a similar memory precision task into the current study design, which allowed us to measure the precision of memory underlying two different reality monitoring decisions (Simons et al., 2008): agency reality monitoring decisions that distinguish actions performed by either the self (i.e., the participant) or an external agent (i.e., the experimenter); and perceptual reality monitoring decisions that distinguish externally perceived and internally imagined experiences. Our novel experiment design bridges previously separate research on reality monitoring and memory precision, and is thereby able to test the extent to which different kinds of reality monitoring decisions may be influenced by the precision of underlying memories.

Our second research question concerns the neurocognitive basis of this possible association. If reality monitoring decisions do indeed depend on memory precision, an experimental manipulation that reduces memory precision should lead to poorer reality monitoring performance. To test this hypothesis, the present study sought to disrupt the functioning of a brain region implicated in long-term memory precision, left angular gyrus (Richter & Cooper et al., 2016), by applying continuous theta burst stimulation (cTBS) (Huang et al., 2005). We predicted stimulation of angular gyrus to reduce memory precision and, in turn, disrupt reality monitoring decisions, compared to stimulation of a control site, the vertex.

One further prediction concerns the possible role of angular gyrus, and its neighbouring posterior parietal regions, in processing self-referential memories (Bonnici et al., 2018; Rugg & King, 2018; Weniger et al., 2009; Yazar et al., 2012, 2014, 2017; c.f. Drowos et al., 2010). If angular gyrus does indeed process personal memories preferentially, stimulation of these regions may disrupt retrieval of self-generated experiences, including both actions performed by the rememberer themselves and internally imagined experiences. We therefore predicted stimulation of angular gyrus to disrupt retrieval of self-referential and imagined experiences disproportionately, relative to actions performed by other people or previous perception of the outside world, ultimately resulting in greater externalisation bias compared to control site stimulation.

In summary, we aimed to answer two key questions stemming from the Source Monitoring Framework: (1) to what extent do reality monitoring decisions depend on the precision with which contextual details of memories are retrieved? and (2) does functioning of angular gyrus represent a neurocognitive basis of this possible dependency? Experiments 1 and 2 investigated whether or not variability in memory precision affects agency and perceptual reality monitoring decisions, respectively. Experiments 3 and 4 investigated whether stimulation of angular gyrus reduces memory precision, compared to control site stimulation, and whether the relatively reduced memory precision disrupts reality monitoring decision making processes. We made three predictions: (1) variability in memory precision will influence reality monitoring performance, (2) cTBS at angular gyrus will reduce memory precision and, in turn, disrupt reality monitoring decisions relative to control site stimulation, and (3) this effect of brain stimulation will disproportionately affect retrieval of self-referential and imagined experiences, relative to externally generated experiences, resulting in a greater magnitude of externalisation bias.

## Methods

### Sample Size Estimation

A priori power analyses were conducted using the ‘pwr.f2.test’ R function to estimate the number of participants with which medium-size effects might be detected *f* = 0.25, *α* = 0.05, *1-β* = 0.8). The analyses were conducted for the following regression models: a model with one outcome variable (i.e., reality monitoring accuracy) and two predictors (i.e., memory precision and reality monitoring conditions) in each behavioural experiment, and a model with an additional predictor (i.e., stimulation site) in each brain stimulation experiment. They revealed that 32 and 37 participants were needed in each behavioural and brain stimulation experiment, respectively. The numbers were increased to 48 participants for each experiment to counter-balance the order of stimulation site in the brain stimulation experiments, and to ensure that the main statistical analyses were comparable across behavioural and brain stimulation experiments *f* = 0.25, *α* = 0.05, *1-β* = 0.95). The number of participants in the current experiments is considerably larger than those in previous studies involving behavioural manipulations and brain stimulation (e.g., Nilakantan et al., 2017; Yazar et al., 2014).

### Participants

Forty-eight participants between 18-35 years of age were included in the analysis for each of the four experiments (192 participants in total). All participants reported normal or corrected-to-normal vision and hearing, and no history of psychiatric or neurological conditions. They provided informed consent in accordance with the procedure approved by the University of Cambridge Human Biology Research Ethics Committee.

Four additional participants withdrew: one person reported discomfort after administration of a single motor threshold pulse, another one reported nausea after four motor threshold pulses, one reported discomfort about 10s after administration of cTBS, and one reported pain 5 minutes after (but not during) administration of cTBS. Sessions were terminated immediately when participants reported any discomfort, nausea, or pain, following approved safety protocols. No participants reported subsequent side effects.

One further volunteer withdrew from Experiment 4, explaining that they could not follow the instructions for the ‘imagined’ reality monitoring condition because of self-reported inability to imagine visuospatial information or ‘aphantasia’ (Zeman et al., 2015).

### Stimuli

Participants were presented with visual displays, each of which consisted of an image of a real-world object overlaid on a faint patterned background. Both the images of objects and the backgrounds were obtained from a real world object stimuli bank from Brady and colleagues (2008) (https://bradylab.ucsd.edu/stimuli.html), and Google image searches. The pairing between objects and backgrounds were randomised. The target location of each object was allocated randomly at a location around a circle between 1-360° angle. The displays were allocated randomly to reality monitoring conditions (i.e., ‘self’, ‘experimenter’, ‘imagined’, and ‘perceived’) and stimulation sites (i.e., angular gyrus and vertex). Displays were identical between participants, but the order of display presentation within each study-test cycle was randomised for each participant. A total of 360 displays were used in the current study.

### Procedures

#### Experiment 1: Agency Reality Monitoring

Participants were asked to complete a practice task followed by 7 study-test cycles. Each cycle consisted of 24 displays. Half of these displays were presented in the ‘self’ condition, where the participants themselves moved an object, and the other half were presented in the ‘experimenter’ condition.

In each study phase trial, a cue was presented (lasting 1s) indicating whether the participant or the experimenter was to move the object that was subsequently presented at the centre of the computer screen. In the ‘self’ condition, participants were asked to hold down the spacebar on a keyboard to move the object progressively to its target location. In the ‘experimenter’ condition, participants were asked to watch the object move progressively to its target location as the experimenter held down the ‘Q’ key. Both the participant and the experimenter had 3s to move the object to its target location. When the object reached its target location, participants were asked to think about what the object was, where its target location was, and whether they themselves or the experimenter had moved the object (3s).

In each test phase trial, a studied object was presented at a random location around an invisible circle. Participants were asked to first recreate the target location of each object by holding down the left and right arrow keys, which moved the object continuously anti-clockwise and clockwise around the circle, respectively. Participants were also asked to indicate whether the object had been moved by the participant themselves or the experimenter during the preceding study phase by pressing the ‘S’ and ‘E’ keys, respectively. The trials were subject paced, terminating if participants did not respond after 9s. Text in the centre of the computer screen reminded participants what each key represented. This text reminder turned red to let participants know when 3s was left before the trial expired.

#### Experiment 2: Perceptual Reality Monitoring

Participants were asked to complete a practice task followed by 10 study-test cycles. Each cycle consisted of 18 displays, half of which were in the ‘imagined’ condition, and the other half were in the ‘perceived’ condition. We used fewer displays for each study-test cycle in Experiment 2 (i.e., 18 displays) relative to Experiment 1 (i.e., 24 displays) to prevent floor effects that emerged in pilot testing of Experiment 2.

In each study phase trial, an object was presented at the centre of the computer screen (3s). In the ‘imagined’ condition, the object was replaced by a black cross, and participants were asked to imagine the object moving progressively to the target location as the cross moved progressively to the target location (1s). In the ‘perceived’ condition, participants were asked to watch the object move progressively to its target location (1s). When the cross or the object reached its target location, participants were asked to think about what the object was, where its target location was, and whether the movement of the object had been imagined or perceived (3s).

Each test phase trial in Experiment 2 was identical to those in Experiment 1 with participants recreating the target location of each object, except that participants were asked to press the ‘I’ and ‘P’ keys to indicate whether the movement of the object had been imagined or perceived during the preceding study phase, respectively (9s). See Figure 1 for an overview of the procedures in Experiments 1 and 2.

**Figure 1.**
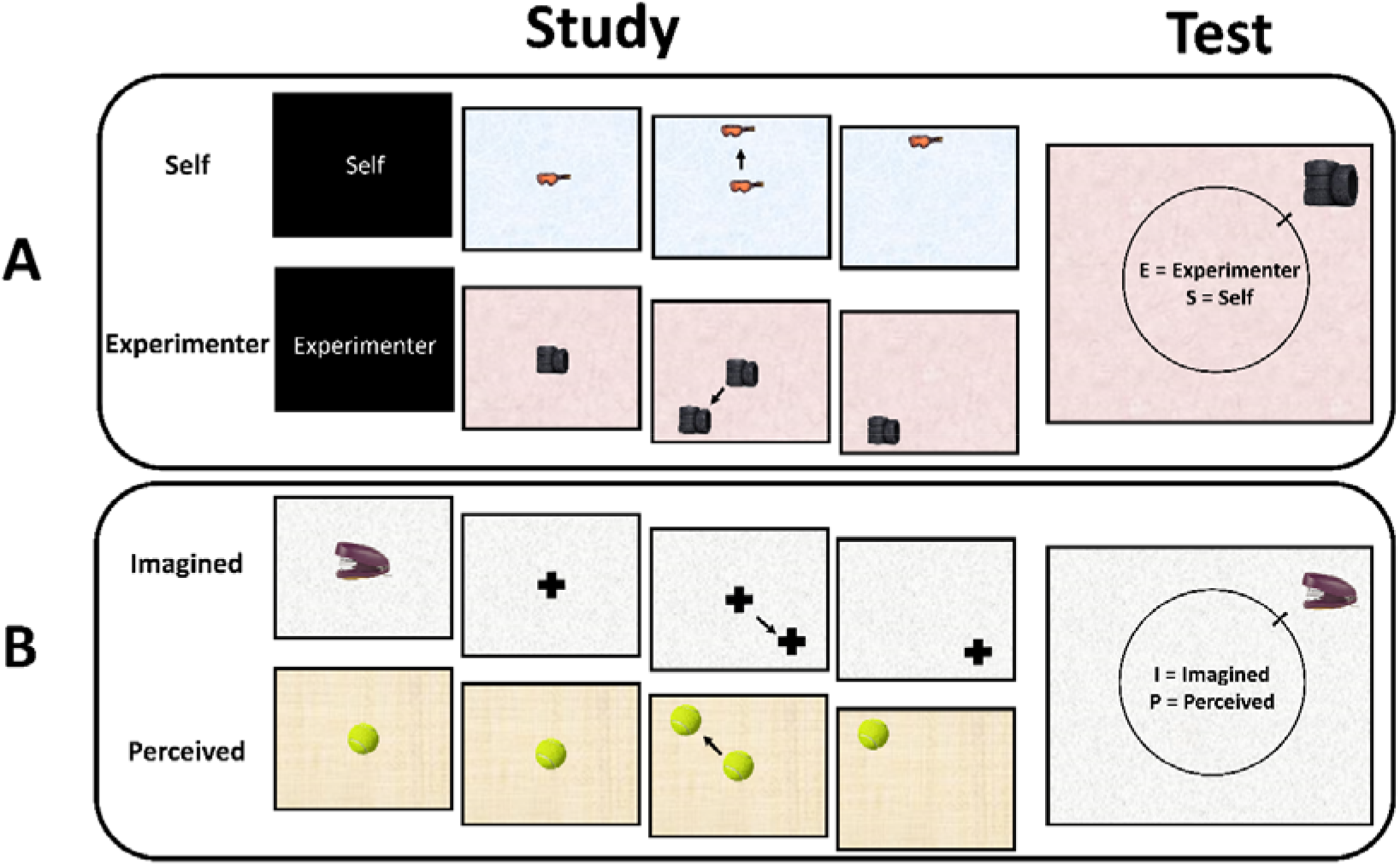
Reality Monitoring Tasks. In the agency reality monitoring task (A), participants were presented with a cue, which indicated whether the participant or the experimenter was to move the following object (1s). Participants were asked to either move the object progressively from the centre of the screen to its target location (i.e., a random location around a circle), or to watch the experimenter move the object to its target location (3s). In the later test phase, participants were asked to recreate the target location and indicate whether the participants themselves or the experimenter had moved the object (9s). In the perceptual reality monitoring task (B), participants were presented an object at the centre of the computer screen (3s). In the ‘imagined’ reality monitoring condition, the object was replaced by a black cross and participants were asked to imagine the object moving progressively to the target location as the black cross progressively moved to the target location. In the ‘perceived’ reality monitoring condition, participants were asked to watch the object move progressively to its target location (1s). In the later test phase, participants were asked to recreate the object location and indicate whether the movement of the object had been imagined or perceived (9s).

#### Experiments 3 and 4: Brain Stimulation

The procedures in Experiments 3 and 4 were identical to those in Experiments 1 and 2, respectively, except each participant received cTBS at angular gyrus and the control stimulation site (i.e., vertex) on separate stimulation sessions. The order of stimulation site was counterbalanced. To prevent the effect of stimulation accumulating between sessions, stimulation sessions were separated by at least 48 hours (and no longer than a week).

Experiments 3 and 4 sought to control for the possibility that cTBS at angular gyrus might disrupt either perception or recognition of objects during the study phase and confound our measure of participants’ later retrieval performance. To control for this possibility, we asked participants to make a semantic decision at the end of each study phase trial, in which they had to successfully perceive and recognise each object in order to decide whether the object would fit inside a shoebox or not. Participants were asked to indicate ‘yes’ and ‘no’ by pressing ‘Y’ and ‘N’ keys, respectively (within 3s or until a response was made).

### Brain Stimulation Methods

The target stimulation site was left angular gyrus, centred on coordinates identified previously in which average peak hemodynamic activity tracked memory precision (Montreal Neurological Institute coordinates: −54, −54, 33) (Richter, Cooper, et al., 2016). The control stimulation site was the probabilistic anatomical vertex (MNI coordinates: 0, −15, 74) (Okamoto et al., 2004). To identify these stimulation sites on each participant’s scalp, half of the participants were registered to their respective T1-weighted structural magnetic resonance head scan, whereas the other half of the participants, for whom no structural scan was available, were registered to the average MNI template head scan provided by Brainsight 2.3.3. The registration method made no significant difference to the main results. See Figure 2 for an illustration of the target and control stimulation sites.

**Figure 2.**
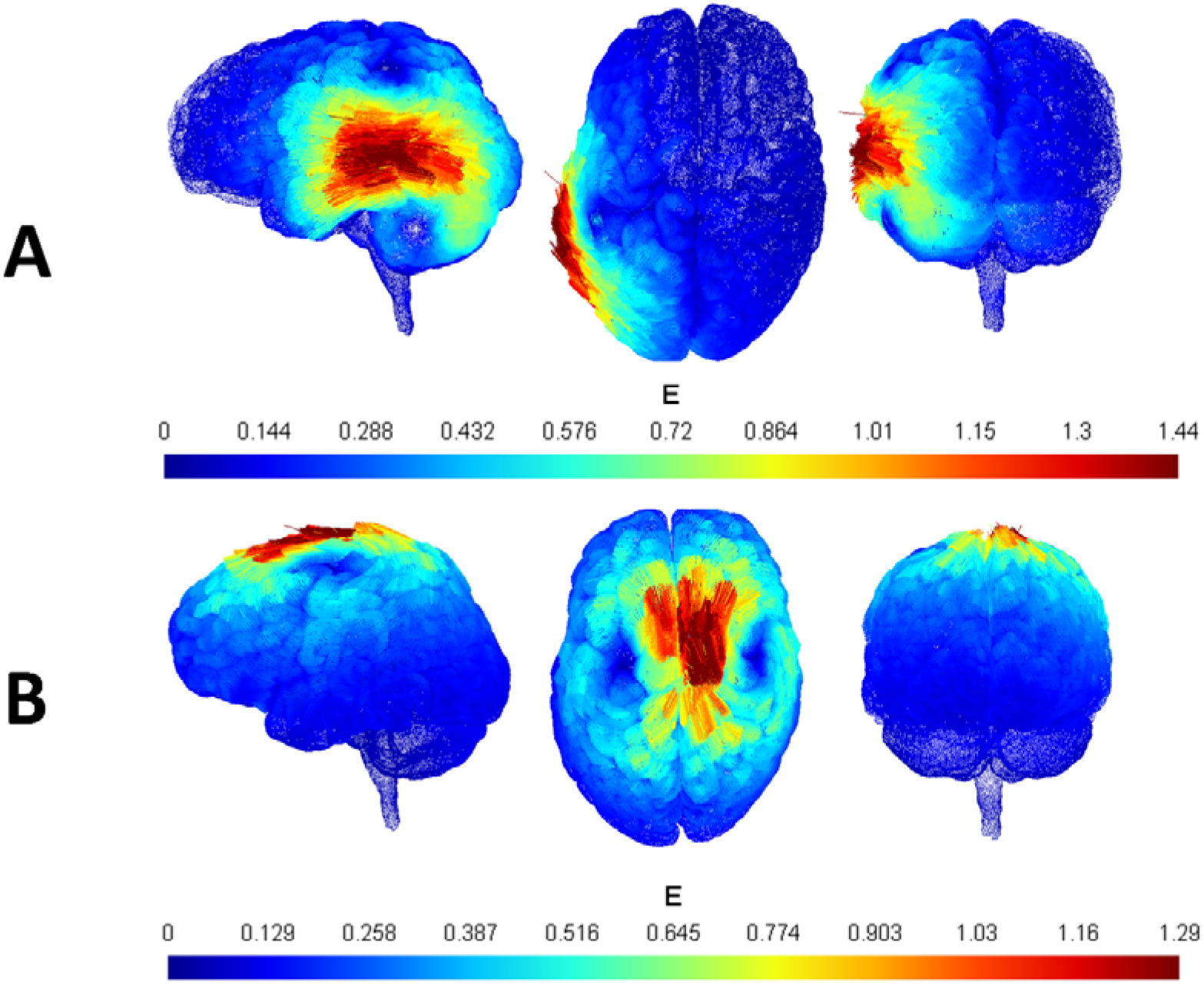
Brain Stimulation Sites. Participants in Experiments 3 and 4 received brain stimulation at angular gyrus (MNI: −54, −54, 33) (A), and the control stimulation site vertex (MNI: 0, −15, 74) (B). Stimulation of each site was separated into two sessions on different days, and the order of stimulation site was counterbalanced. The colours in the figure represent simulations of electric field ‘E’ generated by a 70mm figure eight coil when delivering a single electromagnetic pulse to a template brain, generated using the software ‘SimNIBS’.

To induce electrical currents in participants’ brains, we used the Magstim Rapid 2 stimulator connected to a 70mm figure-eight coil. The resting motor threshold of each participant was estimated by identifying the minimum intensity required to elicit a motor response in each participant’s right index finger or thumb. To this end, we used the TMS Motor Threshold Assessment Tool 2.0 (https://www.clinicalresearcher.org/software.htm). Participants who did not show any motor response were administered the default resting motor threshold, which is 70% of the maximum stimulator output. A standard cTBS protocol was used, where three pulses at 50Hz were delivered every 200ms for 40 seconds at 70% of resting motor threshold. All TMS procedures in the current study were in line with safety guidelines (Rossi et al., 2009, 2021). The distance between the target stimulation site and the centre of the coil did not exceed 2mm during the 40s cTBS for all participants in both Experiments 3 and 4.

### Analysis Approach

In all experiments, we excluded trials with less than 0.5s reaction time or no responses (852 trials were excluded out of 33,408 trials, or 0.256% of all trials). Reality monitoring accuracy was measured on a binary scale: ‘1’ for correct and ‘0’ for incorrect responses. To estimate memory precision, we first calculated the angular error (in degrees) between the target location angle and participants’ recreation of the target location angle. The angular error was then subtracted from the maximum possible angular error (i.e., 180°), so that higher values represented higher memory precision. To prevent random guesses from confounding the estimate of memory precision, trials with less than or equal to 5% likelihood of originating from successful recollection were excluded (see Bays et al., 2009; Richter, Cooper et al., 2016; Schneegans & Bays, 2016). Memory precision was scaled using the ‘scale’ function in the ‘standardize’ package in the programming language R, to reduce the likelihood of discrepancies between the scales of memory precision and other variables, such as reality monitoring accuracy, reality monitoring condition, and stimulation site.

Excluding random guess trials varied the number of remaining observations between reality monitoring conditions, stimulation sites, and participants, potentially confounding within-subject level analysis. To enable within-subject analyses, we matched the number of observations in each condition by randomly sampling, with replacements, trials with more than 95% likelihood of successful recollection. The number of sampled data points were the maximum number of trials in each reality monitoring condition: 84 for each agency reality monitoring condition, and 90 in each perceptual reality monitoring condition. To reduce resampling bias, all analyses were permutated in increments of 100 permutations, until the average standard error of all model outputs across resampling approached 0. Outputs from these permutations of regression models, including generalised eta squared values, were averaged across all permutations. These average permutated model outputs are reported in the results section below.

## Results

### Reduced Memory Precision is Associated with Externalisation Bias

Analyses in behavioural Experiments 1 and 2 first sought to assess whether or not participants exhibited externalisation bias, by testing if reality monitoring performance tended to be reduced in the internal compared to external reality monitoring conditions. Repeated measures ANOVAs testing for an effect of reality monitoring condition on reality monitoring accuracy revealed significant effects in both Experiment 1 (*F*(1,47) = 39.75, *ges* = 0.02, *p* < 0.001) and Experiment 2 (*F*(1,47) = 12.74, *ges* = 0.09, *p* = 0.001). In both experiments, reality monitoring accuracy was significantly reduced in the internal compared to external reality monitoring conditions, consistent with externalisation bias (see Table 1 for descriptive statistics).

**Table 1.**
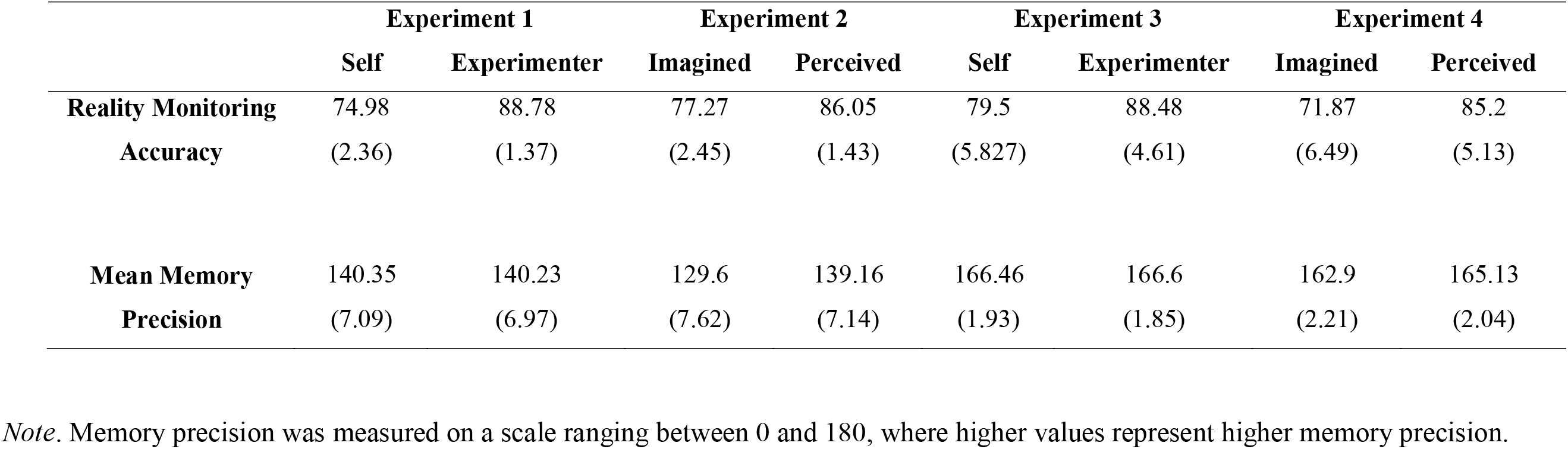
Mean (Standard Error) Reality Monitoring Accuracy (%) and Memory Precision in Experiments 1 to 4.

To assess whether the observed externalisation bias might be linked to reduced memory precision, subsequent analyses tested whether memory precision tended to be reduced in the internal relative to the external reality monitoring conditions. Repeated measures ANOVAs testing for an effect of reality monitoring condition on memory precision did not reveal an effect in Experiment 1 (*F*(1,47) = 0.88, *ges* = 0.001, *p* = 0.41) but a significant effect was observed in Experiment 2 (*F*(1,47) = 19.59, *ges* = 0.008, *p* < 0.001), where memory precision tended to be reduced when participants imagined objects moving to the target location, rather than watching while the objects moved.

We then sought to test directly whether memory precision in each reality monitoring condition (i.e., ‘self’, ‘experimenter’, ‘imagined’, and ‘perceived’ conditions) tended to be associated with reality monitoring performance. To estimate these possible associations, we computed mixed effects logistic regression models within each reality monitoring condition, where the outcome variable was reality monitoring accuracy, the predictor was memory precision, and the random effects variable was participants. These regression models revealed significant positive associations in the ‘self’, ‘imagined’, and ‘perceived’ reality monitoring conditions (*β* ≥ 2.73, *R*^2^ ≥ 0.3, *p* ≥ 0.014), whereas no such association was observed in the ‘experimenter’ reality monitoring condition (*β* = 1.92, *R^2^* = 0.32, *p* = 0.27).

We further analysed these associations to test if, for example, memory precision tends to be relatively reduced, participants are more likely to misattribute self-referential memories to other people. If participants did indeed exhibit this possible ‘It had to be you’ effect in the agency reality monitoring experiment, the observed association between memory precision and reality monitoring performance would be disproportionately greater in the ‘self’ condition compared to the ‘experimenter’ condition. Repeated measures ANOVA comparing the association between ‘self’ and ‘experimenter’ condition did indeed reveal a significant effect of reality monitoring condition (*F*(1,47) = 9.69, *ges* = 0.12, *p* = 0.003), where the association was greater in the internal ‘self’ condition (*M* = 4.36, *SE* = 0.51) compared to the external ‘experimenter’ condition (*M* = 1.91, *SE* = 0.42). We also sought to test if, conversely, in instances where memory precision is disproportionately greater, participants are more likely to misattribute imagined experiences to be real. This possible ‘It had to be real’ effect would correspond to a disproportionately smaller association in the ‘imagined’ condition compared to the ‘perceived’ condition. Repeated measures ANOVA comparing the association between ‘imagined’ and ‘perceived’ conditions did indeed reveal a significant effect of reality monitoring condition (*F*(1,47) = 5.48, *ges* = 0.05, *p* = 0.024), where the association was smaller in the internal ‘imagined’ condition (*M* = 2.77, *SE* = 0.43) compared to the external ‘perceived’ condition (*M*= 4.25, *SE* = 0.44). An a-posteriori mixed ANOVA comparing these effects observed in Experiments 1 and 2 revealed a significant cross-over interaction between reality monitoring conditions (i.e., internal and external conditions), and experiments (*F*(3,141) = 6.1, *ges* = 0.09, *p* = 0.006).

### Stimulation of Angular Gyrus Does Not Reduce Control Semantic Task Performance

Turning to Experiments 3 and 4, the first set of analyses aimed to test whether cTBS at angular gyrus might have reduced participants’ ability to perceive or recognise objects, relative to control stimulation. If there was indeed an effect of stimulation on semantic control task performance, we sought to assess whether reality monitoring conditions were affected disproportionately. We therefore compared participants’ performance on the control semantic judgement task between stimulation sites and reality monitoring conditions. In both Experiments 3 and 4, ANOVAs comparing participants’ accuracy on the control semantic task between stimulation sites and reality monitoring conditions did not reveal any significant effect of stimulation site (*F*(1,47) ≤ 0.45, *ges* ≤ 0.002, *p* ≥ 0.507) or reality monitoring condition (*F*(1,47) ≤ 0.95, *ges* ≤ 0.003, *p* ≥ 0.335). No significant interaction was observed between stimulation site and reality monitoring condition (*F*(1,47) ≤ 1.04, *ges* ≤ 0.004, *p* ≥ 0.313).

### Stimulation of Angular Gyrus Does Not Reduce Memory Precision or Reality Monitoring Accuracy

The main analyses in both Experiments 3 and 4 aimed to test whether cTBS at angular gyrus might reduce memory precision and reality monitoring performance, compared to control site stimulation. Looking first across reality monitoring conditions, repeated-measures ANOVAs revealed no significant main effect of stimulation site on memory precision (*F*(1,47) ≤ 0.79, *ges* ≤ 0.004, *p* ≥ 0.438) or reality monitoring accuracy (*F*(1,47) ≥= 0.27, *ges* ≥= 0.001, *p* >= 0.658).

We then sought to test the prediction that cTBS at angular gyrus might selectively reduce memory precision and reality monitoring accuracy in the internal relative to external reality monitoring conditions. Repeated measures ANOVAs testing possible interactions between stimulation site and reality monitoring condition did not, however, reveal an interaction effect on memory precision (*F*(1,47) ≥= 0.36, *ges* ≥= 0.001, *p* >= 0.439) or reality monitoring accuracy (*F*(1,47) ≥= 4.99, *ges* ≥= 0.002, *p* >= 0.074). Similarly, pair-wise comparisons testing possible effects of stimulation site for each reality monitoring condition revealed no significant effects on memory precision (t(1,47) ≤ 1.15, *d* ≤ 0.24, *p* ≥ 0.29, *BF* ≤ 0.36) or reality monitoring accuracy (t(1,47) ≤ 1.69, *d* ≤ 0.35, *p* ≥ 0.13, *BF* ≤ 0.89).

### Stimulation of Angular Gyrus Reduces the Association Between Memory Precision and Self-Referential Reality Monitoring Accuracy

The next analyses tested whether cTBS at angular gyrus reduced the associations observed between memory precision and reality monitoring accuracy. Looking first across reality monitoring conditions, repeated measures ANOVAs testing for an effect of stimulation site on the observed association between memory precision and reality monitoring accuracy did not reveal a significant effect (*F*(1,47) ≤ 1.29, *ges* ≤ significantly reduced the association in the; 0.001, *p* ≥ 0.081).

To test our directional prediction that cTBS at angular gyrus might reduce disproportionately the association between memory precision and reality monitoring accuracy in the internal reality monitoring conditions compared to external conditions, we computed separate mixed effects regression models for each reality monitoring condition at each stimulation site. These regression models revealed significant positive associations between memory precision and reality monitoring accuracy in all reality monitoring conditions at both stimulation sites (*β* ≥ 1.98, *R^2^* ≥ 0.19, *p* ≤ 0.002), consistent with the results of Experiments 1 and 2. We then sought to test whether the main effect of cTBS at angular gyrus reduced disproportionately the observed association in the internal reality monitoring conditions, compared to the external conditions. Repeated measures ANOVAs testing whether an interaction between reality monitoring conditions and stimulation sites affected these associations did indeed reveal a significant interaction in the agency reality monitoring task (*F*(1,47) = 14.46, *ges* = 0.07, *p* < 0.001), whereas no such interaction was observed in the perceptual reality monitoring task (*F*(1,47) = 1.95, *ges* = 0.007, *p =* 0.17). Follow-up pair-wise comparisons in the agency reality monitoring task revealed cTBS at angular gyrus to have significantly reduced the association in the ‘self’ reality monitoring condition (t(1,47) = 4.25, *d* = 0.87, *p* < 0.001) but not in the ‘experimenter’ condition (t(1,47) = 1.36, *d* = 0.31, *p* = 0.18). A mixed ANOVA directly comparing the effects observed in agency and perceptual reality monitoring tasks revealed a significant three-way interaction between reality monitoring condition, stimulation site, and experiment (*F*(1,94) = 4.33, *ges* = 0.01, *p =* 0.04). See Figure 3 for an illustration of this three-way interaction.

**Figure 3.**
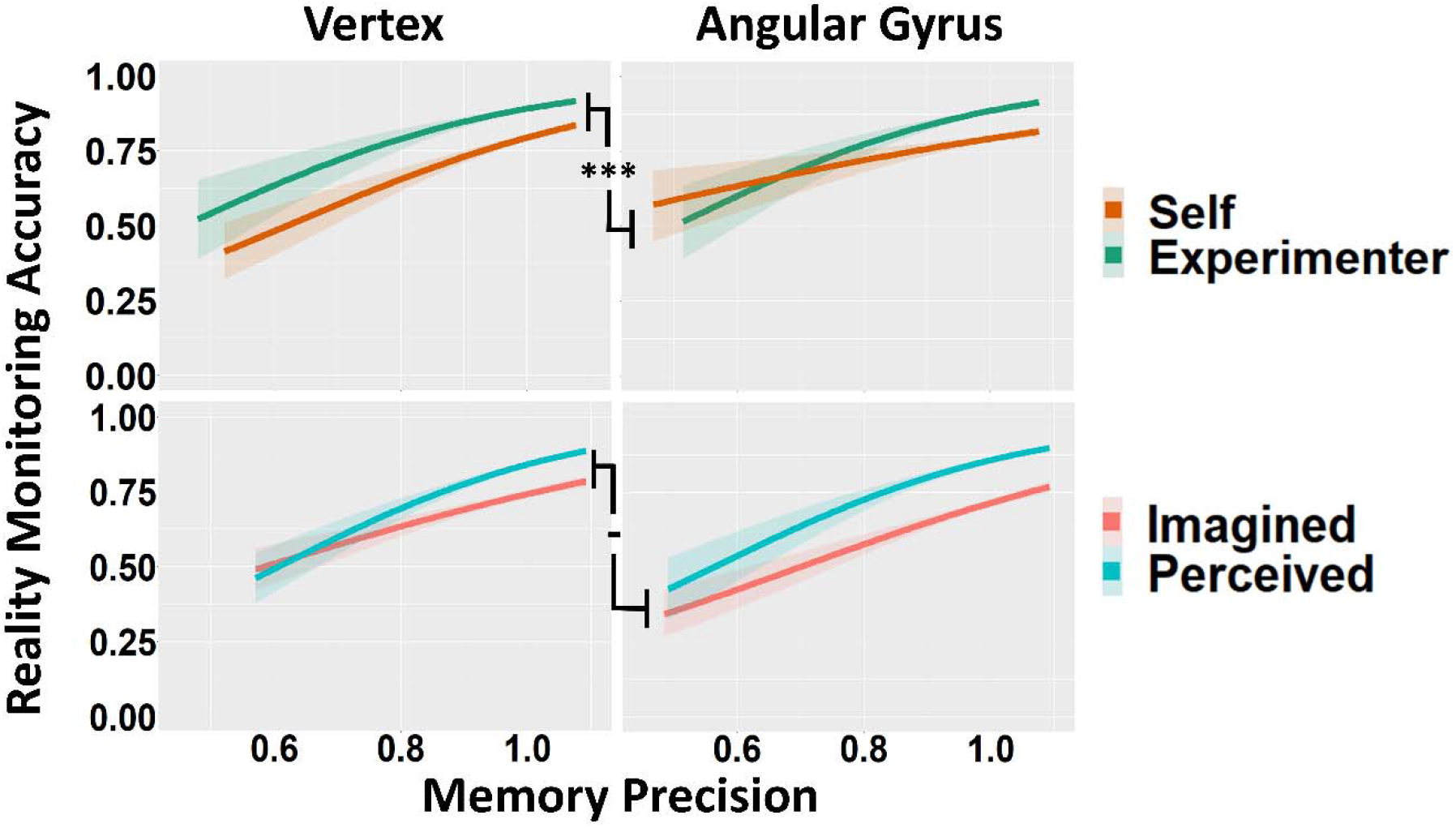
Associations Between Memory Precision and Reality Monitoring Accuracy. Plots in the top and bottom rows of the figure represent performance in agency and perceptual reality monitoring tasks, respectively. Plots on the left and right sides of the figure represent task performance under vertex and angular gyrus stimulation, respectively. The lines represent mean regression coefficients, where the outcome variable is reality monitoring accuracy and the predictor is memory precision. The shaded areas represent standard error of the respective mean regression coefficient. ‘-’:*p* > 0.05, ‘***’:*p* < 0.001.

## Discussion

The current study aimed to test a key hypothesis stemming from the Source Monitoring Framework, that reality monitoring decisions may depend, at least partly, on the quality of the underlying mnemonic representation (Johnson et al., 1993). We sought to test this prediction by combining a memory precision task with agency and perceptual reality monitoring tasks in behavioural Experiments 1 and 2, respectively. Both experiments revealed relatively reduced memory precision to be associated with lower reality monitoring performance, substantiating the Source Monitoring view. To investigate a possible neurocognitive basis of the observed behavioural association, Experiments 3 and 4 aimed to reduce memory precision by applying cTBS at angular gyrus and test if the relatively reduced memory precision disrupts reality monitoring performance. Stimulation at angular gyrus, relative to control site stimulation, had no effect on overall memory precision or reality monitoring performance. However, when participants recalled actions they had performed as opposed to actions carried out by another person, stimulation of angular gyrus selectively reduced the association between memory precision and self-referential reality monitoring decisions, compared to the control site stimulation. In light of these findings, we discuss below how angular gyrus may be important for the ability to imbue remembered experiences with a sense of self-agency, enabling a key component of ‘autonoetic consciousness’ that characterises episodic memory (Tulving, 1985).

The current behavioural findings suggest that reduced episodic memory precision might obscure the self-referential features that characterise a mnemonic representation as the rememberer’s personal experience. Previous studies have found that participants expect to experience a sense of self-agency when remembering their past actions (Johnson et al., 1981), but if they cannot remember who performed a task, they tend to misattribute their own actions to another person, rather than the other way around, yielding the ‘It had to be you’ effect (Johnson & Raye, 1981). This effect or ‘agency externalisation bias’ has also been observed in previous studies involving patients with schizophrenia, where the magnitude of the bias was greater among the patients than healthy controls, suggesting that these patients’ ability to imbue remembered experiences with a sense of self-agency may be reduced compared to healthy people (Bentall et al., 1991; Seal et al., 1997; Waters et al., 2006; Woodward et al., 2007). Patients may, for example, misattribute their self-generated actions to another person, or to other external sources, such as voices of people they cannot see, or shadows of people they cannot touch physically.

Whereas a sense of self-agency is particularly important in self-referential reality monitoring decision making, visuoperceptual features of a remembered experience may be more relevant in perceptual reality monitoring decisions. If, for example, patients with schizophrenia experience vivid multisensory hallucinations, those imaginary experiences may be misattributed as real, at least partly because the vivid and rich multisensory imagined stimuli resemble real experiences (Mondino et al., 2019). Even healthy people may exhibit instances of perceptual externalisation bias where, for example, they report having seen objects moving when, in fact, they had only imagined the movement of objects, especially if they retrieve precise visuoperceptual details of a similar but different experience (Richter, 2020). In line with such previous findings, participants in the current study might have been more likely to misattribute imagined experiences as real, rather than the other way around, if the internally generated experiences were unusually vivid or precise, or if the participants mistakenly incorporated into their memory rich visuoperceptual details of a similar previous experience.

These links between reduced memory precision and externalisation biases may help to resolve mixed findings in previous studies (Alderson-Day et al., 2019; Allen et al., 2006, 2006; Collignon et al., 2005; Garrison et al., 2017; Larøi et al., 2004; Moseley et al., 2021). Those previous studies sought to test whether healthy people with greater proneness to hallucinations may exhibit cognitive profiles often associated with psychosis, such as externalisation biases. However, the tasks used typically involved categorical responses to measure mental experiences that might have obscured variability in the qualitative characteristics of the mental representations tested. The present study observed relatively robust links between memory precision and externalisation biases, perhaps because the continuous measure of memory precision provided additional sensitivity that was able to reveal a link between reality monitoring decisions and the qualities of underlying mnemonic representations.

Turning to the possible neurocognitive basis of memory precision and reality monitoring, stimulation of angular gyrus did not reduce the overall precision of recollection or reality monitoring decision making. This observation suggests the possibility that when the functioning of angular gyrus is disrupted, other functionally connected regions such as hippocampus may step in and support memory performance. If this is the case, enhancing the functional connection between hippocampus and angular gyrus might be expected to increase memory precision. Consistent with this possibility, a previous study stimulated the functional connection between hippocampus and posterior parietal regions including angular gyrus, and observed the stimulation to increase memory precision, substantiating the notion that interconnected regions may step in to support recollection (Nilakantan et al., 2017). In typical situations, however, it is likely that these functionally connected brain regions work together to support precise recollection (Cooper & Ritchey, 2019).

One must always be cautious, however, in attempting to interpret a null result as in the present data, as it might simply reflect insufficient power or another technical deficiency such as failure to stimulate the correct underlying brain region. Because of these possible reasons, it might be that the current brain stimulation experiments were unable to observe the stimulation of angular gyrus to reduce memory precision. Note, however, that the sample size in the present experiments was supported by power calculations and was considerably larger than in many previous brain stimulation studies of memory (e.g., Nilakantan et al., 2017; Yazar et al., 2014). Furthermore, the present data revealed a significant effect of angular gyrus stimulation on the relationship between memory precision and self-referential reality monitoring performance (discussed below), which suggests that these experiments were capable of revealing such significant effects if they did exist.

The observation that stimulation of angular gyrus selectively reduced the link between memory precision and self-referential reality monitoring decisions is consistent with proposals that functioning of angular gyrus may contribute to the self-referential quality of memories (Bonnici et al., 2018; Rugg & King, 2018; Weniger et al., 2009; Yazar et al., 2012, 2014, 2017). This self-referential mnemonic function may involve integrating multiple modalities of retrieved features, such as visual and auditory details (Yazar et al., 2017), within an egocentric, first-person perspective framework (Bonnici et al., 2018), in order to reconstruct rich and subjectively vivid mnemonic representations that are imbued with a sense of self-agency (Simons et al., 2021; Zou & Kwok, 2021).

The rememberer may then evaluate whether this sense of self-agency resembles self-referential characteristics often expected of personal memories, enabling the reality monitoring decision making that is thought to be supported by functioning of anterior medial prefrontal cortex (Simons et al., 2017). Although angular gyrus and anterior medial prefrontal cortex may support episodic memory precision and self-referential reality monitoring decision making, respectively, it is likely to be through their interaction that a holistic subjective experience of remembering may arise (Simons et al., 2021). This ability to project rememberers themselves into subjective mental experiences characterises the ‘autonoetic consciousness’ that defines episodic memory (Tulving, 1985).

On the basis of the evidence considered above, it seems reasonable to have expected that, at least in the current Experiment 3, stimulation of angular gyrus might have disrupted participants’ ability to experience autonoetic consciousness and, in turn, reduced self-referential reality monitoring performance. Performance in that condition remained intact, however, suggesting that reality monitoring decisions may not solely depend on just one quality of mnemonic representations; the rememberer might instead consider other qualities of the underlying mnemonic representation, such that performance may be maintained even in situations where one of the ways to experience autonoetic consciousness is disrupted. If, for example, the precise visuoperceptual location of an object had been forgotten, the rememberer might instead retrieve other visuoperceptual features such as the colour or orientation of the object and, in turn, consider whether these precisely remembered features resemble characteristics often expected of real and imagined experiences. Such adaptations may allow the rememberer to adequately distinguish such experiences in most situations, resulting in the measurement of intact reality monitoring performance.

In summary, the current study observed relatively reduced memory precision to be associated with lower reality monitoring performance, substantiating the key notion from the Source Monitoring Framework: reality monitoring ability may depend, at least in part, on the quality of the underlying mnemonic representation. We also observed participants to misattribute their past actions to other people, especially when the underlying memory precision tended to be reduced, consistent with the ‘It had to be you’ effect (Johnson & Raye, 1981). Conversely, participants were more likely to misattribute imagined experiences as real, rather than the other way around, if imagined experiences were remembered with unusually precise visuoperceptual details that meant they resembled veridical events. Turning to the neurocognitive basis of these observed links between memory precision and reality monitoring ability, angular gyrus may be important for the ability to imbue remembered experiences with a sense of self-agency, enabling the autonoetic consciousness that characterises episodic memory.

## Acknowledgements

This work was supported by a James S McDonnell Foundation Scholar Award to JSS. It was completed within the University of Cambridge Behavioural and Clinical Neuroscience Institute, funded by a joint award from the UK Medical Research Council and the Wellcome Trust. For the purpose of open access, the author has applied a CC BY public copyright licence to any Author Accepted Manuscript version arising from this submission. We are very grateful to Zoe Kourtzi and Andrew Welchman for use of their TMS system, and to Lukas Schaffner and Mihaela Taranu for TMS assistance. We thank Paul Bays for helpful advice.

